# Self-organized morphogenesis of a human neural tube *in vitro* by geometric constraints

**DOI:** 10.1101/2021.07.24.453659

**Authors:** Eyal Karzbrun, Aimal H. Khankhel, Heitor C. Megale, Stella M. K. Glasauer, Yofiel Wyle, George Britton, Aryeh Warmflash, Kenneth S. Kosik, Eric D. Siggia, Boris I. Shraiman, Sebastian J. Streichan

## Abstract

Understanding how human embryos develop their shape is a fundamental question in physics of life with strong medical implications. However, it is challenging to study the dynamics of organ formation in humans. Animals differ from humans in key aspects, and in particular in the development of the nervous system. Conventional organoids are quantitatively unreproducible and exhibit highly variable morphology. Here we present a morphologically reproducible and scalable approach for studying human organogenesis in a dish, which is compatible with live imaging. We achieve this by precisely controlling cell fate pattern formation in 2D stem cell sheets, while allowing for self-organization of tissue shape in 3D. Upon triggering neural pattern formation, the initially flat stem cell sheet undergoes folding morphogenesis and self-organizes into a millimeter long anatomically accurate model of the neural tube, covered by epidermis. We find that neural and epidermal human tissues are necessary and sufficient for folding morphogenesis in the absence of mesoderm activity. Furthermore, we find that molecular inhibition of tissue contractility leads to defects similar to neural tube closure defects, consistent with *in vivo* studies. Finally, we discover that neural tube shape, including the number and location of hinge points, depends on neural tissue size. This suggests that neural tube morphology along the anterior posterior axis depends on neural plate geometry in addition to molecular gradients. Our approach provides a new path to study human organ morphogenesis in health and disease.

## Main

Brain and spinal cord development begins with the folding of the embryonic neural tissue into a tube (*1*, *2*) (Fig. 1a). Defects in neural folding are one of the most common birth defects affecting 1:1000 pregnancies, which can result in severe disabilities and lethality shortly after birth (*3*–*5*). This highlights the need for understanding the cellular and tissue scale processes which drive neural tube folding in humans (*6*–*8*). To address the limited accessibility in human embryos, there has been an accelerated development of 3D human stem cell cultures (organoids) capable of recapitulating selected aspects of human organ formation (*9*–*11*). Organoids have greatly advanced our understanding of cell fate decisions during organ formation, yet they yield variable and anatomically incorrect tissue shapes and cell-fate patterns (*12*) (Fig. 1b). Additional organs-on-chip systems have been developed to apply stem-cells into scalable, controlled and functional tissues (*13*). However, the integration of stem-cells in complex microfluidic environments over-constrains their shape, and does not allow for self-organization as in embryonic development. Thus, new approaches are required in order to study human organ morphogenesis.

**Fig. 1.**
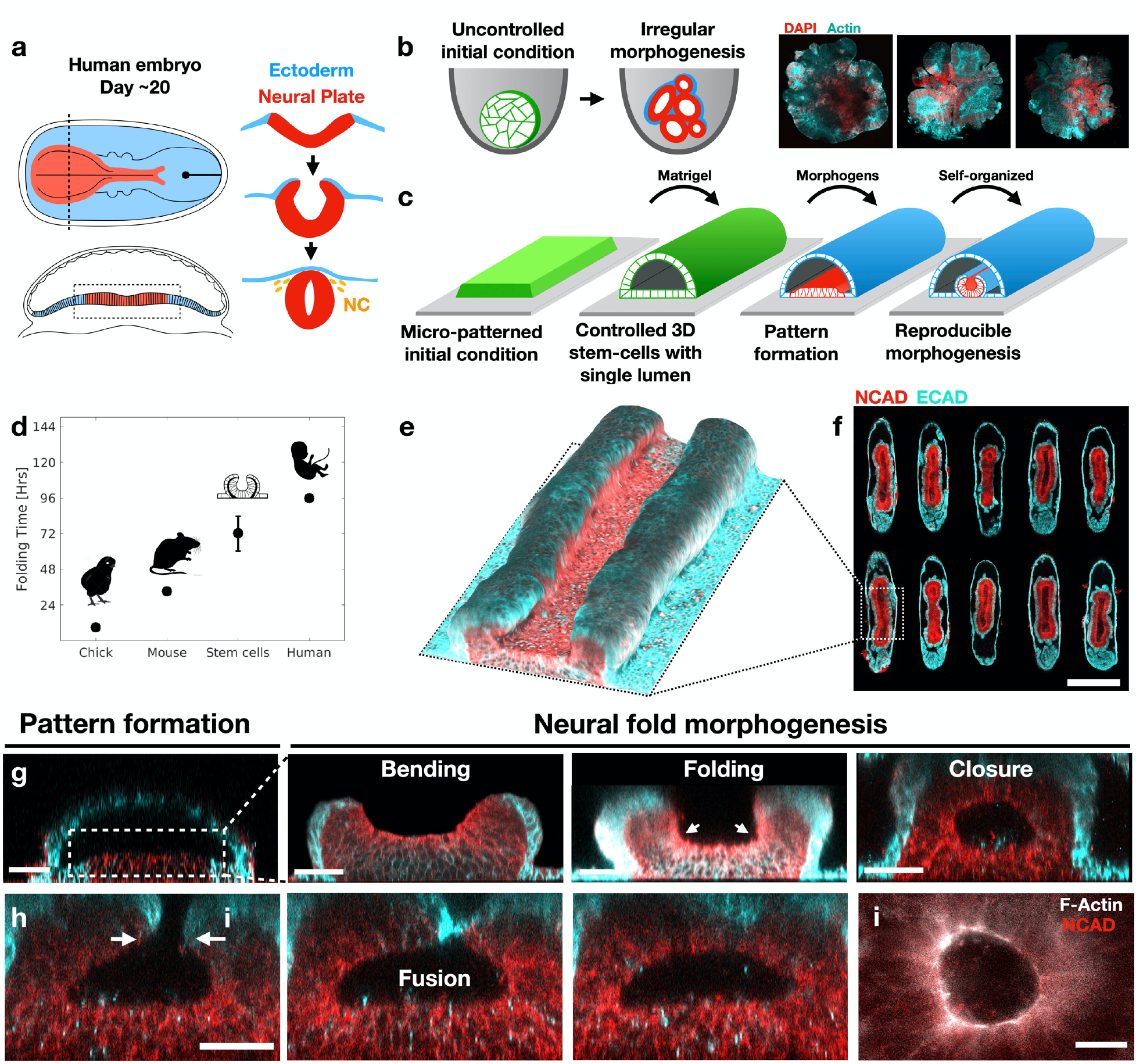
A reproducible human stem-cell model of neural-tube morphogenesis. (a) Scheme of neural fold morphogenesis in a human embryo. Neural plate shown in red, Non neural ectoderm in blue, and neural crest (NC) in yellow. (b) Uncontrolled initial conditions result in irregular morphogenesis observed in conventional organoids. (c) Controlled initial conditions lead to reproducible cell-fate pattern formation and morphogenesis. Scheme showing key steps in protocol. See Methods for detail. (d) Neural folding duration in the stem-cell system compared to folding time in chick (*41*), mouse (*42*), humans (*17*). Time is measured between neural plate formation to first fusion of the neural folds. (e) 3D-reconstruction of a stem-cell derived ~1mm long neural tube. (f) Horizontal section through a micropatterned array of stem-cell derived neural tubes with reproducible morphology. (g) Stages of neural folding in the stem cell system. (h) Neural closure is mediated by fusion of the non-neural ectoderm layer. (i) An actin ring is observed during neural closure. Scale bars 500μm (f), 50μm (g,h), 25μm (i).

### A reproducible stem cell system to study human organ morphogenesis

Here, we present a new experimental system that allows us to study the dynamics of organ morphogenesis using human stem-cells (Fig. 1c). Our system is reproducible, scalable, and compatible with live imaging and genetic manipulations. The experimental design is inspired by the embryonic development of organs. Organs develop from a primary embryonic tissue, which is precisely controlled in all physical aspects including cell number, shape and size. In many cases this primordium is a two-dimensional sheet of polarized cells (epithelium) in contact with a lumen, which forms an isolated biochemical niche. This provides a well-defined physical and biochemical starting point for cell-fate patterning in the 2D epithelium that subsequently triggers 3D organ shape formation. We thus hypothesized that controlling the initial physical conditions in which organoids develop, and realizing a 3D epithelial culture with a single lumen will enable us to reproducibly recapitulate aspects of human organ morphogenesis.

We solve the reproducibility challenge of stem-cell cultures by applying surface micropatterning. In this way we are able to precisely control the initial size, shape and cell number of 2D stem-cell cultures (Fig. S1). We then apply Matrigel to trigger a robust transition from the constrained 2D culture into a 3D pluripotent epithelium that wraps around a single large lumen. The 3D stem-cell culture shape and size are controlled by the micropattern geometry and forms at >95% success rate (Fig. S2–3). Importantly, the 3D tissue maintains pluripotency, and can thus give rise to many cell types of the human body (Fig. S4). Moreover, the lumen is physically and chemically isolated from the environment, thus mirroring the *in vivo* situation (Fig. S5). As in embryonic development, the self-organization of stem cells can be guided by exposure to morphogens. In this manuscript we focus on modeling neural tube formation by applying morphogens involved in early neurodevelopment. We further demonstrate the capacity of our system to generate a human amnion and forebrain organoids, thus establishing a broadly applicable system for studying human organ morphogenesis (Fig. S6).

To model neural tube morphogenesis, we expose the 3D stem cell culture to a combination of morphogens which is thought to be involved in early neural development (*14*–*16*). We first apply a neural induction media containing TGFβ inhibitor (SB-431542), followed by exposure to bone morphogenetic protein 4 (BMP4) (Fig 1c, S1). In response to the morphogens, the system exhibits self-organized folding morphogenesis which takes place over three days (Fig. 1d). The *in vitro* folding period is similar to neural folding in human embryos, which takes place over four days between the appearance of the neural plate and the first fusion of the neural folds (*17*). The result is a tube-shaped neural tissue covered with surface ectoderm, which recapitulates multiple anatomical features of the embryonic neural tube (Fig. 1e). The process is highly reproducible and occurs with over 90% success rate in multiple cell lines (Fig. 1f, S7). Notably, fold morphogenesis is not observed in 2D cultures, and requires a 3D tissue surrounding a lumen as an initial condition (Fig. S8).

### Stem-cells recapitulate hallmarks of *in vivo* neurulation

Remarkably, *in vitro* folding morphogenesis follows the sequence of *in vivo* neurulation: neural plate formation and thickening, bending, folding, and closure (Figs. 1g, S9). The columnar neural epithelia dimensions, 215±15μm width x 70±5μm thickness, are comparable to human neural plate in Carnegie stage 8 (Fig. S10) (*17*, *18*). Furthermore, live imaging reveals that neural cells undergo interkinetic nuclear motion typical of neuroepithelia (*19*). Neural bending is concentrated at two focal points, which are formed at the intersection between the uprising neural folds and the glass adhered tissue. These are reminiscent of lateral hinge points which form during neural tube development *in vivo*. Finally, the neural closure occurs via a zippering motion in which the non-neural ectoderm makes the first contact (Fig. 1h), and an actin ring is observed at the closing edge (Fig. 1i), as during *in vivo* neural closure (*20*, *21*).

Another hallmark of neural tube development is the formation of a tissue bilayer composed of a neural layer and surface ectoderm (Fig. 2a). An immunostaining study reveals that the stem cell system indeed forms a bilayer of anterior neural tissue (NCAD, PAX6, OTX2, Fig. 2b-e), and non-neural surface ectoderm (ECAD, KTN8, TFAP2*α*, Fig. 2f-g). We further observe a population of neural crest cells at the neural-ectoderm interface, which coincides with their place of origin *in vivo* (SOX10, PAX7, Fig. 2h-i). Interestingly, the neural-ectoderm interface is enriched in fibronectin and focal adhesions suggesting that the bi-layer is formed by basal adhesion of the two cell types (Fig. 2j-k). Fibronectin is observed exclusively at the neural-ectoderm interface, whereas the ectoderm-glass interface is enriched with collagen (Fig. S11). Similar ECM composition has also been observed *in vivo*, where the dorsal neural-ectoderm interface is enriched with fibronectin, whereas the ectoderm-mesoderm interface is enriched with collagen (*22*, *23*). Overall, these findings suggest that the *in vitro* model system recapitulates key aspects of *in vivo* neurulation, in terms of morphology and timing, cell fate patterns, extracellular matrix, and cellular behaviors.

**Fig. 2.**
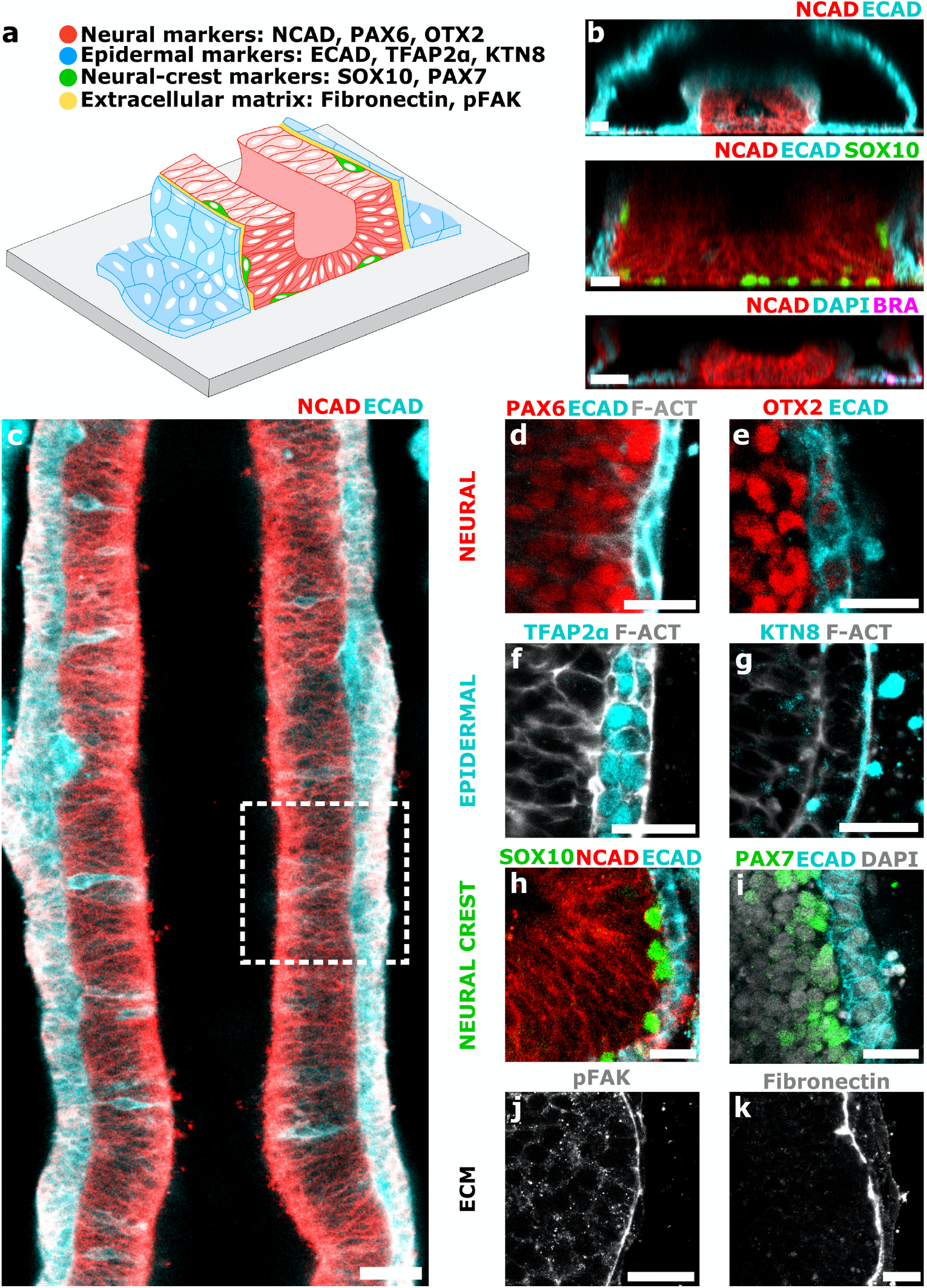
Formation of a neural-ectoderm bilayer. (a) Scheme of a neural-ectoderm bilayer formed during stem-cell fold morphogenesis. The neural-ectoderm interface is enriched with extracellular matrix, and neural crest cells. (b) Vertical and (c-k) horizontal sections through the stem cell derived neural tube. Immunostaining with mesoderm marker Brachyury (BRA) reveals no mesoderm tissue is involved in neural folding. Samples additionally immunostained with anterior neural markers PAX6 and OTX2 (d,e); epidermal markers TFAP2α and Kaertain8 (KTN8) (f,g); Neural crest markers SOX10 and PAX7(h,i); Focal adhesion marker phosphorylated focal adhesion kinase (pFAK), and extracellular matrix marker fibronectin (j,k). Scale bars 50μm (b,c) 25μm (d-k).

### Epidermal and neural tissues are necessary and sufficient for folding

We next apply our system to study which tissues contribute to neural folding. Over the past decades work in animal models has been instrumental in revealing the cellular behaviors and signaling pathways which underlie neural tube closure. In particular, it was shown that cellular behaviors in the neural ectoderm, epidermis, and mesoderm all play a role in neural folding (*2*). Our system now allows us to address the question of which tissues are required for neural tube folding in a human background. The *in vitro* system allows to systemically control tissue composition through the choice of small molecules in the culture media (Fig. S6). In this way we are able to dissect out the tissues which are necessary and sufficient for tube morphogenesis. Immunostaining with mesoderm marker Brachyury shows that there is no mesoderm tissue present during folding morphogenesis (Fig. 2b). Mesoderm fates are suppressed by TGFβ inhibition, and only appear if the period of TGFβ inhibition is decreased (Fig. S12). Thus, our data suggest that mesoderm is not required for folding morphogenesis *in vitro*. We next examine whether the non-neural ectoderm is required for folding. For this purpose, we employed prolonged neural induction without BMP4. This results in a homogenous neural tissue that lacks epidermal markers, and does not exhibit folding morphogenesis (Fig. S6). Finally, we observe that addition of BMP4 without neural induction lead to mesoderm and ectoderm cell fates, but lack neural tissue and does not exhibit folding (Fig. S12B). We thus conclude that epidermal and neural tissues are necessary and sufficient to drive folding morphogenesis. Our data suggest that mesoderm activity is not required for neural folding and closure, in contrast to previous studies in animal models (*24*, *25*).

### Towards modeling neural tube defects

Neural tube defects (NTD) are a severe and prevalent birth defect. Mouse and additional animal models have successfully recapitulated genetic, cellular and morphological aspects of NTD (*5*). In addition to animal model systems stem-cell culture methods offer a complementary approach to study human-specific aspects of NTD (*26*, *27*). Here we wanted to test if our stem-cell system can be used to model NTD. As a first step towards this goal, we tested whether we could recapitulate the defects observed in neural tube defects using small molecular inhibitors. For this purpose we focused on Shroom3-Rho kinase (ROCK) signaling, which is associated with human NTDs (*28*, *29*). Shroom3 is upregulated in the neural tissue, and localizes ROCK to actin fibers in the apical surface of the neural plate (*2*, *30*–*32*). To perturb ROCK/Shroom3 signaling and apical neural contractility we apply a highly specific molecular inhibitor of ROCK (Y-27632) (Fig. 3a). ROCK inhibition results in severe folding defects similar to the open book morphology observed in embryonic cranial NTDs (Fig. 3b-d). ROCK inhibited samples exhibit a thick and flat neural tissue, which lacks neural hinges and is significantly less curved than control samples (Fig. 3e). The neural apical cell areas and total apical tissue area are significantly larger in ROCK inhibited samples, whereas epidermal cell areas do not exhibit a significant change (Fig. 3c, d, f-g). Furthermore, Shroom3 and F-actin localization to the apical neural surface are disrupted in response to ROCK inhibition, indicating that our assay perturbs shroom signaling, disrupts apical actomyosin assembly, and prevents apical contraction of the neural tissue (Fig. 3h-k). In contrast, actin localization to the basal surface of epidermal/neural tissues, as well as the epidermal apical surface, is not significantly perturbed. Taken together, these data demonstrate that interference with ROCK/Shroom3 signaling leads to perturbation of apical neural contractility and results in folding defects. This indicates that neural contractility is required for folding morphogenesis. Taken together this suggests that our *in vitro* model system is suitable for studying how abnormalities in signaling pathways lead to neural tube defects in a human genetic context.

**Fig. 3.**
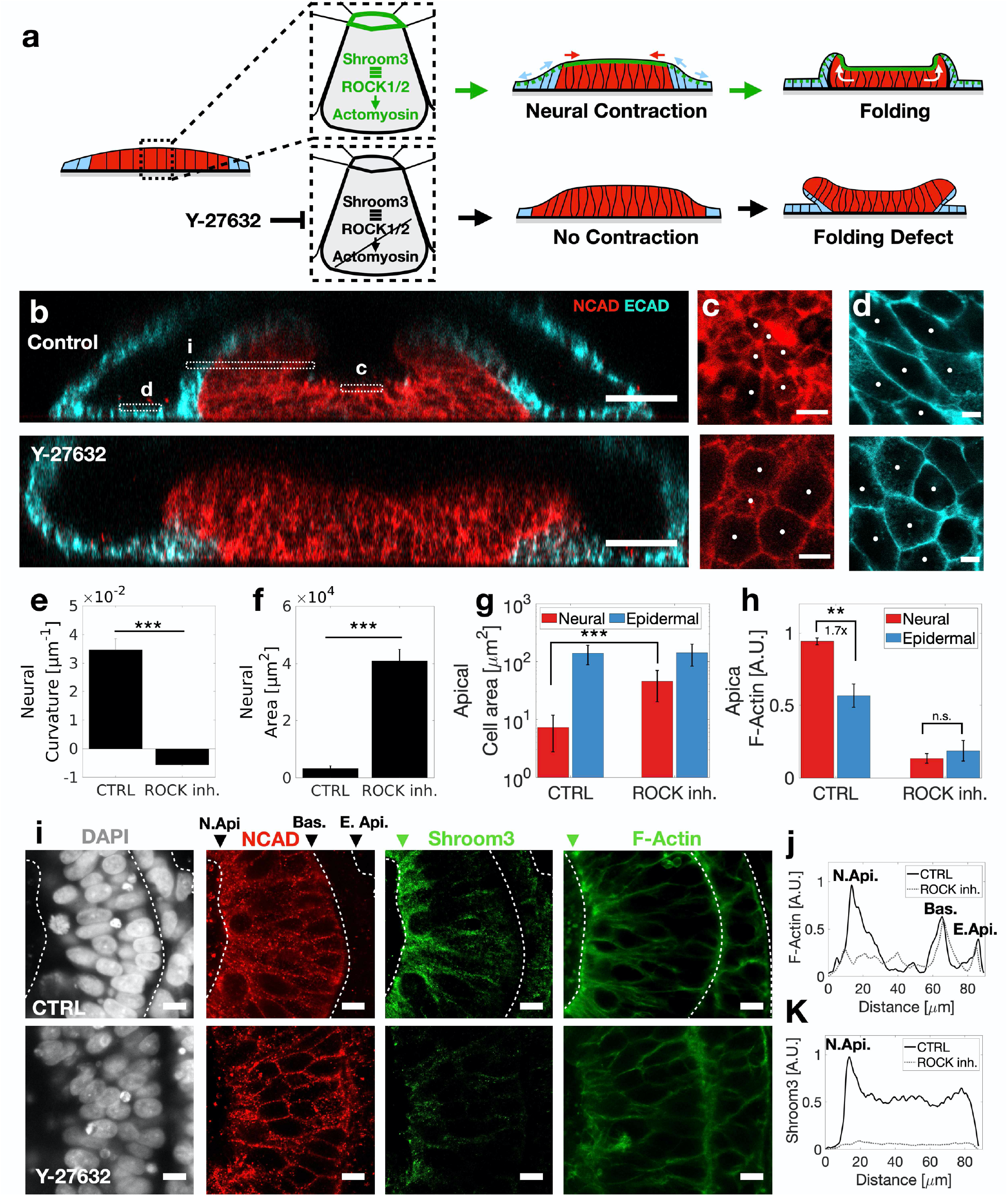
Modeling neural tube defects. (a) Schematic of neural contractility and folding in the presence of active ROCK-Shroom3 signaling. A folding defect is observed in response to inhibition of ROCK signaling using the inhibitor Y-27632. (b) Vertical sections showing neural fold defect in response to ROCK inhibition. (c) High magnification of neural (red) and (d) epidermal (cyan) apical cell borders in control and ROCK inhibited samples. (e) Neural curvature is significantly lower in ROCK inhibited samples than in control (CTRL) samples (***, p<0.005). (f) Total neural tissue apical area in ROCK inhibited samples at 72hpb is significantly higher than in CTRL. (g) Neural apical cell area significantly increases in ROCK inhibited samples, whereas epidermal cell area does not change. (h) F-actin intensity is higher in apical neural tissue than in apical epidermal tissue (**, p<0.01). The apical F-Actin gradient is lost in response to ROCK inhibition. (g) Horizonal sections show Shroom3 and F-actin localization to neural apical surface (N. Api.). (j) F-Actin and (k) Shroom3 fluorescence intensity across the apical-basal axis in CTRL (solid black line) and ROCK inh. (grey dashed line) samples. F-Actin intensity is highest in the neural apical surface, and is disrupted in response to ROCK inhibition. (J) (K) Scheme showing ROCK inhibition downregulates actin gradient, prevents neural contraction, and leads to folding defect. Data are mean ± S.E.M., N=3 samples(C, F), N=40 (E). Scale bars 50μm (B), 5μm (D, D’), 10μm (G).

### Neural plate size determines neural tube shape

The neural plate width varies along the anterior-posterior (AP) axis, from ~500μm on anterior brain region down to 100-200μm on the posterior end (Fig. 4a) (*33*). Neural fold morphology also varies along the anterior-posterior axis, exhibiting a broad fold with two lateral hinges at the anterior end, and narrow fold with a single medial hinge at the posterior end. The current paradigm is that neural tube morphology is controlled by cell behaviors driven by a gradient of signaling molecules along the AP axis (*34*, *35*). However, to which extent the neural plate size directly controls shape remains unclear.

**Fig. 4.**
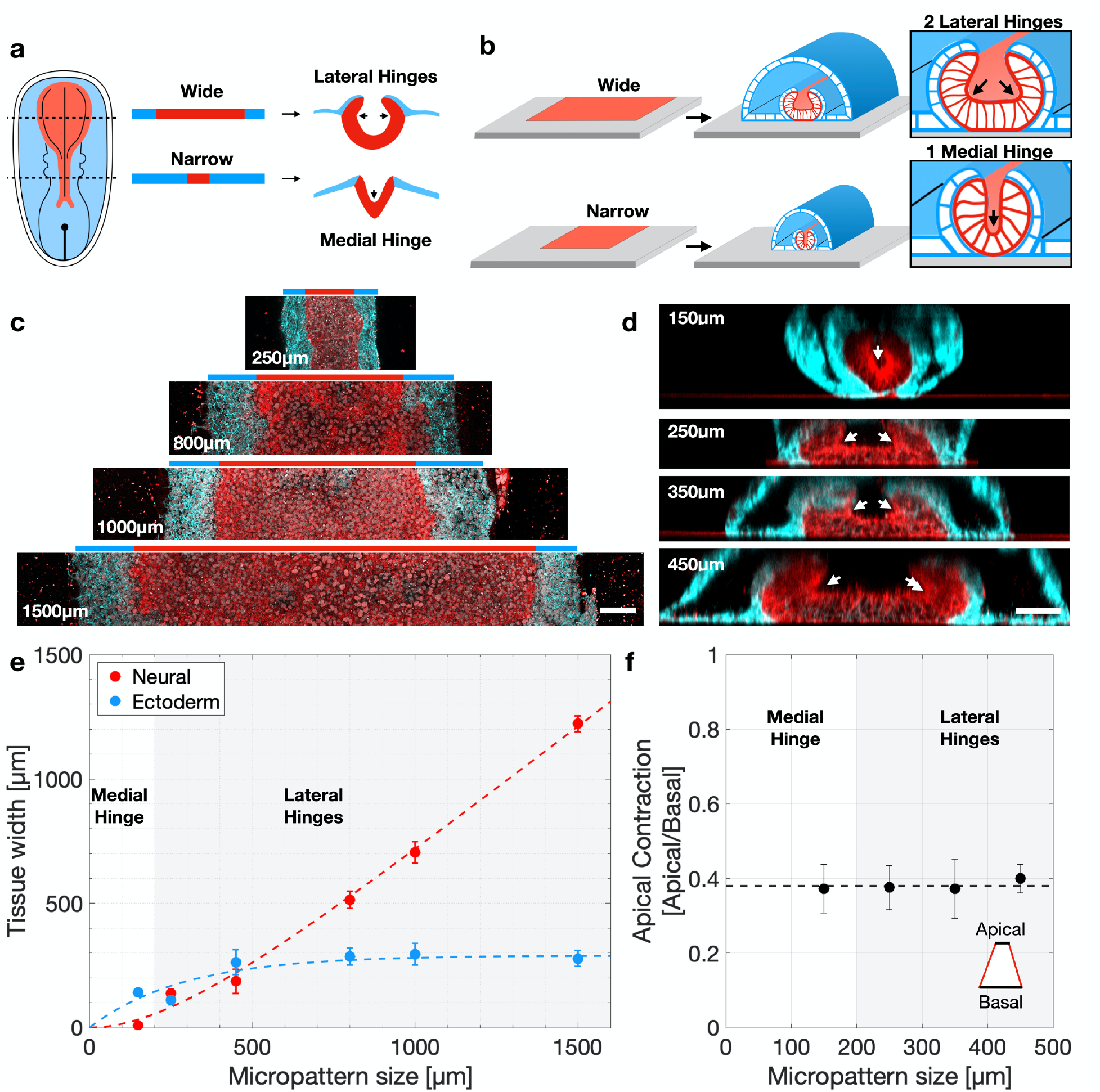
Neural plate size determines neural tube shape. (a) Scheme showing changes in neural plate size and neural tube shape along the anterior posterior axis. (b) Experimental design - micropatterns are created at increasing widths to study the effect of tissue size on fold morphology. (c) Horizonal sections show neural (NCAD, red) and ectoderm (ECAD,cyan) pattern formation at varying micropattern widths. (d) Vertical sections reveal neural fold morphology changes as micropattern width increases. At 150um a single medial hinge is observed, whereas larger micropatterns result in two lateral hinges. (e) Plot of neural and ectoderm tissue width as a function of micropattern size. White area indicates the region in which a single hinge is observed. Grey area indicates the region in which two hinges are observed. (f) Plot of the ratio of apical to basal neural area as a function of micropattern size. The apical-basal ratio is independent of micropattern size and is indicative of apical contractility. Scale bars 100μm (c) and 50μm (d).

We tested how tissue size controls shape by micropatterning rectangular geometries at increasing widths (Fig. 4b). We observe that the width of the neural epithelium scales linearly with micropattern size, whereas the surface ectoderm is restricted to a fixed region of ~150μm at the boundary of the micropattern (Fig. 4c, e). Surprisingly, folding morphology does not simply scale with increasing size (Fig. 4d). Instead, narrow micropatterns (150μm width), exhibit a u-shaped fold with a single central hinge point, characteristic of posterior regions. In contrast, wide micropattern (>150μm width) carry two lateral hinges, characteristic of anterior regions. Notably, neural epithelium apical contractility did not change as a function of tissue size, suggesting that the different morphologies do not arise from changes in cell behaviors (Fig. 4f).

To better understand the conditions for the appearance of hinge points we apply a computational model (SI, Fig. S13). We find that a combination of apical neural contractility and basal adhesion between epidermal and neural tissues are sufficient for neural fold morphogenesis and hinge point formation *in silico*. Furthermore, each hinge point is characterized by a finite size. In cases where the neural tissue size is of order of the hinge point size, a single hinge point appears the center of the neural tissue, whereas for larger tissues two hinge points appears. Overall, our data suggest a new scenario, in which a combination of tissue mechanics, patterning geometry, and cell behaviors determine the final shape of the neural tube.

## Discussion

In summary, we developed a reproducible model system for studying human organogenesis in a dish. Our approach enables us to address a broad range of problems in human development. In particular, we recapitulate key features of *in vivo* neurulation in terms of molecular and cellular composition and tissue morphogenesis. We further demonstrate a reproducible and high-throughput model system which is compatible with live imaging and genetic manipulations, and suitable for a future study of the human genetics which underlie neural tube defects. Surprisingly, our results indicate that neural tube shape, and in particular the number and location of hinge points, is directly controlled by neural plate size. This may contribute to the morphological differences along the anterior-posterior axis. Furthermore, the observation that fold shape does not scale with the neural epithelium size, suggest that neural folds form by forces at the neural-ectoderm interface. Finally, our results suggest that the presence of a lumen is important for folding morphogenesis. Whether the lumen provides essential biochemical or physical cues remains to be determined. In the future, our approach can be applied to study human specific aspects of other organs, and to pave way for synthetic morphogenesis.

## Acknowledgments

S.J.S. acknowledges NIH grant R21 HD099598-0. E.K. acknowledges the Human Frontier Science Program for the long-term postdoctoral fellowship (LT000629/2018-L). A.H.K. acknowledges support of the National Science Foundation Graduate Research Fellowship Program (Grant no. 1650114). EDS acknowledges NSF grant 2013131. S.M.K.G was recipient of an Early Postdoc Mobility grant and a Postdoc Mobility grant from the Swiss National Science Foundation (P2ZHP3_174753 and P400PB_186800). We thank Y. Huang and the Gladstone institute for sharing CTR2 hiPSC line, and Brivanlou lab and the Rockefeller University for sharing RUES2 embryonic and reporter lines. We thank Noah P. Mitchell for sharing computer codes for 3D surface analysis. We thank the Streichan and Kosik lab members, as well as the UCSB stem-cell center and associate director Cassidy Arnold for fruitful discussions and advice. Finally, we thank Alexandra M. Tayar for valuable comments on the manuscript.

## Methods

### Microfabrication and soft lithography

Device fabrication is performed using standard soft-lithography techniques on a four-inch wafer. One layer of photoresist (SU-8 2075, Microchem) is spun onto a silicon wafer at a thickness of 110μm. Photoresist is exposed to ultraviolet light using a mask aligner (Suss MicroTec MA6) and unexposed photoresist is developed away to yield multiple arrays of posts. A Trimethylchlorosilane layer is vapor deposited on the developed wafer to prevent adhesion. A 10:1 ratio of PDMS and its curing agent (SYLGARD 184 A/B, Dow Corning) is poured onto the wafers and cured at 65C overnight. The PDMS layer is then peeled off the silicon mold and individual stamps are cut out using a razor blade for future use.

### Glass micropatterning

Sterile PDMS stamps and 35 mm diameter custom-made glass-bottomed culture dishes are plasma treated for 1 minute on high setting (PDC-32G, Harrick Plasma) to activate both surfaces. Stamps are pressed features-side to the glass surface and held in place. To passivate the glass surface in nonpatterned regions, 0.1mg/mL PLL-g-PEG solution (SuSoS AG, Switzerland) is added to petri dish immediately after securing stamps to glass surface and incubated for 30 minutes. Stamps are then carefully removed and stamped glass dishes are rinsed several times with PBS++. Laminin-521 (STEMCELL Tech.) is added at a dilution of 5μg/mL in PBS++ to incubate overnight at 4°C. The following day, stamped glass dishes are rinsed with PBS++ to remove excess unbound laminin and used within 1-7 days.

### hPSC lines and maintenance

The work reported in this paper was approved by the Human Stem Cell Research Oversight Committee (hSCRO) at University of California Irvine, study # 2018-1072. In this work, we used early passages of hiPSC CTR2#17 (Figs. 1–4) (*36*). The cell line was previously karyotyped as normal (*36*). In addition, we used hiPSC reporter line AICS-0023 developed by the Allen institute (Fig. S2,3,5), and NIH-approved embryonic stem cell line RUES2 NIH approval number NIHhESC-09-0013 (Fig. S9). hPSCs were cultured with mTESR1 media (STEMCELL Tech.) on hESC-qualified Matrigel (Corning) coated dishes. Media was exchanged on a daily basis, and cells were regularly checked for mycoplasma contamination.

### Neural tube morphogenesis protocol

#### Day 1. Seeding onto micropatterns

hPSCs are released from well-plate surfaces using non-enzymatic agitation following manufacturer’s instructions (ReleSR, STEMCELL Tech.). Cells are resuspended as a single-cell suspension at densities of 750K-1M cells/mL in mTeSR1 containing 10μM ROCK inhibitor Y27632 (Abcam). 200μL of cell suspension is then pipetted onto prepatterned dishes and allowed to settle for 15 minutes before adding 1mL of mTeSR1 and allowing cells to settle for 10 additional minutes. Excess media is aspirated, leaving enough liquid to cover patterns and replaced with fresh 2mL of mTeSR until the following day.

#### Day 2. Matrigel addition

mTESR1 media is exchanged with a neural induction media containing Matrigel (4%, v/v). Neural induction media is supplemented with 5μM of TGFβ-inhibitor SB-431542 (Table S1).

#### Day 3-4. Lumen Formation

Dishes are left unperturbed at day 3 to allow transition into 3D stem-cell tissue containing a single lumen.

#### Day 5-9. Exposure to Morphogens

Neural induction media is supplemented with 5ng/mL BMP4 in addition to 5μM of SB-431542. Media is exchanged daily. Cell fates are observed on day 6 and folding is observed during days 7-9.

### Whole-mount immunostaining

All samples were fixed, immunostained and imaged as whole mounts in the culture dish. Samples are fixed in 4% PFA, 1hr at room temperature (RT), washed three times for 15 min in PBS, and permeabilized in 1.5% Triton-PBS over-night (o/n) at 4C. The following day, samples are washed in 0.3% Triton, and blocked for 2hrs RT (10% NGS, 1% BSA, 0.3% Triton-X in PBS). Primary antibodies are used at 1:100-1:200 in blocking solution o/n at 4°C. Next, samples are washed in 0.1% Tween in PBS, and incubated with secondary antibodies 1:500-1:1000 in PBT o/n at 4°C. DAPI and phalloidin are also added at this stage. Finally, samples are washed in PBS for 1hr, and imaged.

### ROCK inhibition

ROCK inhibition is achieved using the small molecule Y-27632 reconstituted in water at a stock concentration of 10mM. ROCK inhibitor was applied at a concentration of 10μM (1:1000 dilution) at day 5, together with BMP, and maintained for 72hrs until the end of the experiment at day 8. Media was changed daily. Control experiments were carried under identical conditions, adding water instead of ROCK inhibitor.

### Imaging

Fluorescence imaging of whole-mount immunostained samples and was carried out using Leica SP8 confocal microscope. Imaging was carried out directly on glass-bottom dishes in which Lumenoids were cultured. Live-imaging was carried out in a custom-made incubator for temperature and CO2 control.

### Three-dimensional surface reconstruction

The 3D image in Fig. 1c was created using an in-house computational pipeline combining (i) ilastik an interactive machine learning for (bio)image analysis (*37*), (ii) ImSAnE an open-source MATLAB toolbox (*38*), and (iii) MeshLab an Open-Source Mesh Processing Tool (*39*). First, a confocal stack is segmented in ilastik. An additional segmentation is carried out to sperate neural and epidermal tissues, based on N/E-Cadherin fluorescence signals. These steps result in 3D stacks with segmentation probability values at each voxel. Then, ImSAnE and MeshLab are used to build a 2D triangular mesh at the apical surface of the tissue. For this purpose, an integral detector using Morphsnakes is used (*40*). Next, ImSAnE is used to project fluorescence intensity values from the original 3D confocal stack onto the 2D mesh.

## Supplementary Figures

**Fig. S1.**
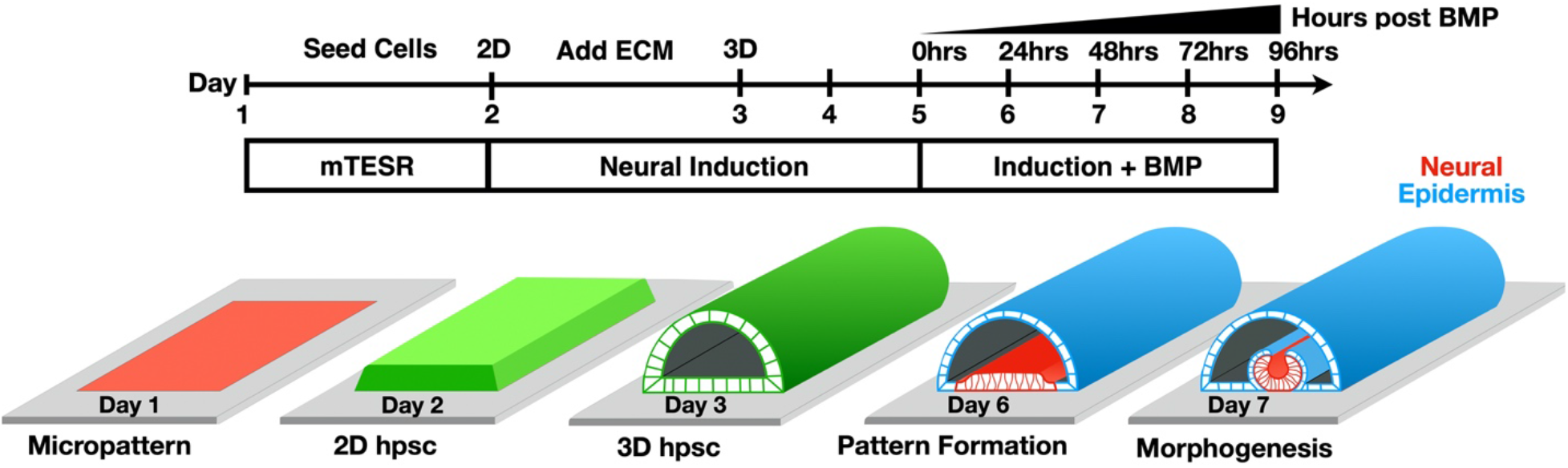
Experimental timeline. Cells are seeded on micropatterns on Day 1. On day 2 neural induction media containing 5μM of TGFβ-inhibitor SB-431542 (Table S1). On Day 3 cells transition into a 3D tissue containing a single lumen. On days 5-9 neural induction media is supplemented with 5ng/mL BMP4 in addition to 5μM of SB-431542. Neural and ectoderm cell fates are observed on day 6 and folding morphogenesis is observed during days 7-9.

**Fig. S2.**
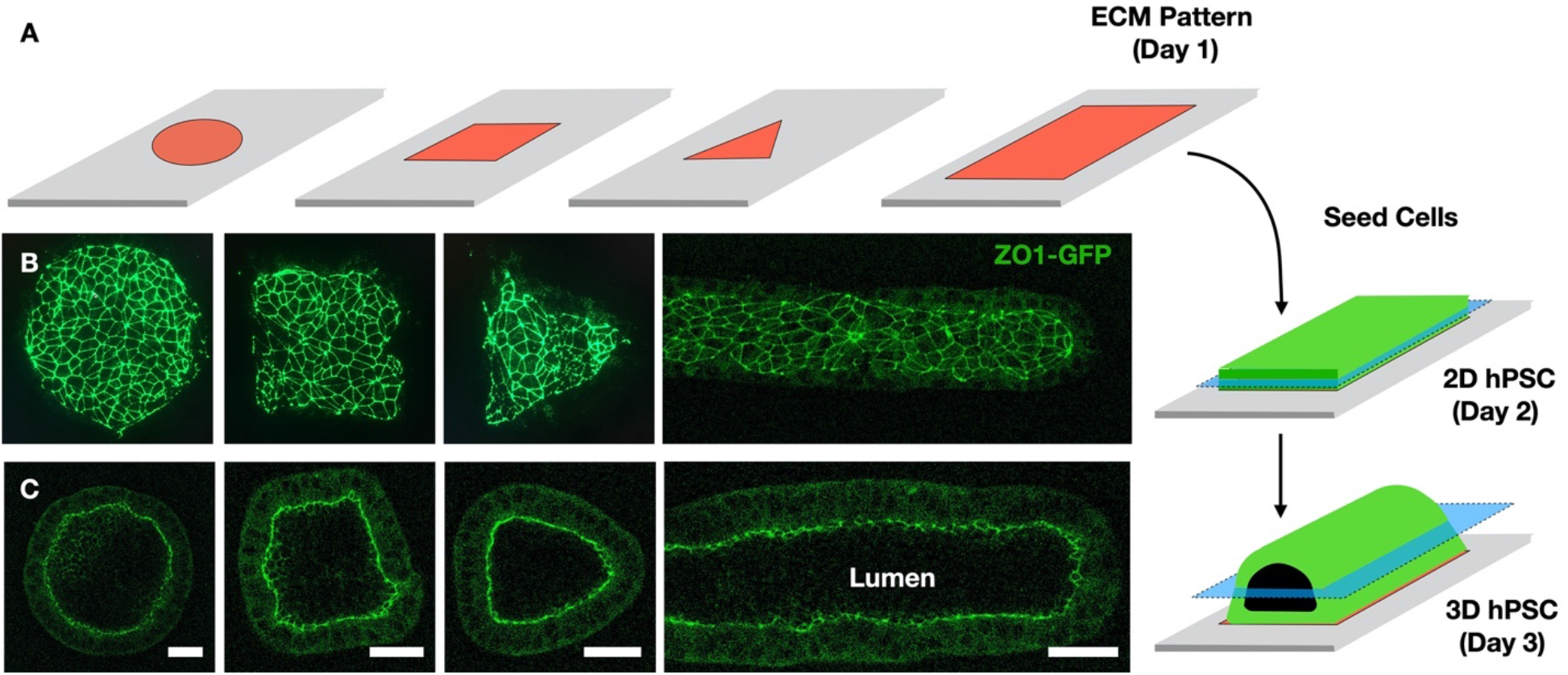
Micropattern geometry controls 3D stem-cell culture shape. (A) Scheme showing shape-controlled ECM pattern deposited on glass surface. (B) Seeding of hPSCs onto micropatterns results in two-dimensional cultures which are restricted to the micropattern geometry. (C) Adding 4% Matrigel to the media results in three-dimensional stem-cell cultures containing a single large lumen. The 3D shape is controlled by the micropattern geometry. Scale bar is 50μm.

**Fig. S3.**
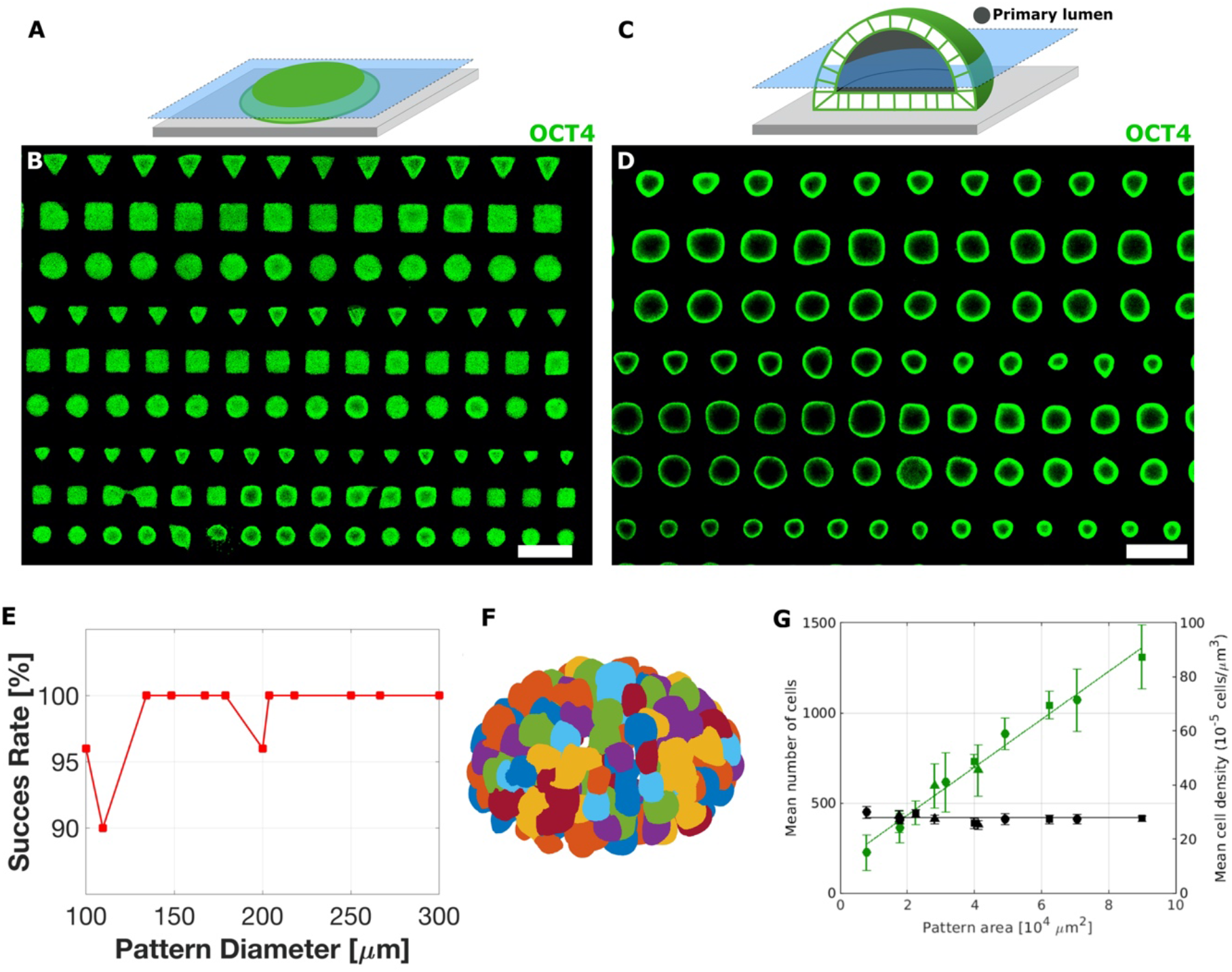
Three-dimensional hPSC with single lumen are reproducible and scalable. (A) Scheme showing micropatterned hPSC and imaging plane. (B) Micropattern array of hPSCs before Matrigel addition (Day 2). The array contains dozens to hundreds of individual colonies. (C) Scheme showing micropatterned 3D hPSC after Matrigel addition. (D) Micropattern array of hPSCs after Matrigel addition (Day 3). (E) 3D hPSC with a single lumen forms with success rate greater than 90%. (F) Segmentation of single nuclei is used to count the number of cell in each sample. Each nucleus is labeled with a different color. (G) Total cell number in each sample scales linearly with pattern area while cell density remains invariant. Total n =300. Scale bar is 500μm.

**Fig. S4.**
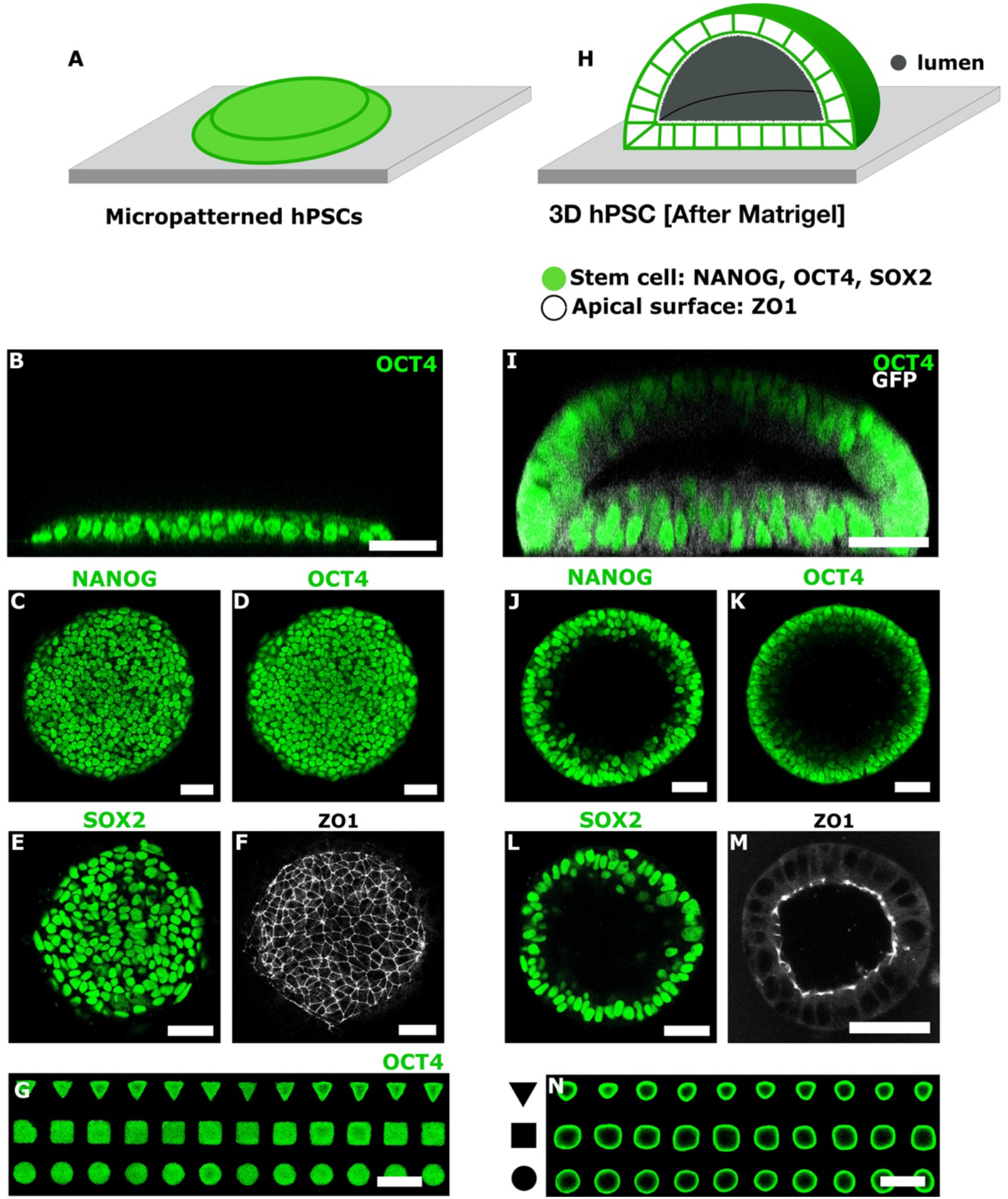
Three-dimensional hPSC form a pluripotent epithelia surrounding a single lumen. (A) Scheme showing micropatterned hPSC. Cells represented in green over grey surface. (B) Vertical section of micropattern hPSC immunostained with pluripotency marker OCT4 (day 2). (C-E) Horizontal sections showing pluripotency markers NANOG, OCT4, and SOX2. (F) Horizontal section near colony top surface showing tight junction protein ZO1. (G) Scheme showing micropatterned 3D hPSC after Matrigel addition. (H) Vertical section of 3D hPSC immunostained with pluripotency marker OCT4 (day 3). (I-K) Horizontal sections showing pluripotency markers NANOG, OCT4, and SOX2. (L) Horizontal section through center of the 3D culture showing tight junction protein ZO1. Scale bar 50 μm.

**Fig. S5.**
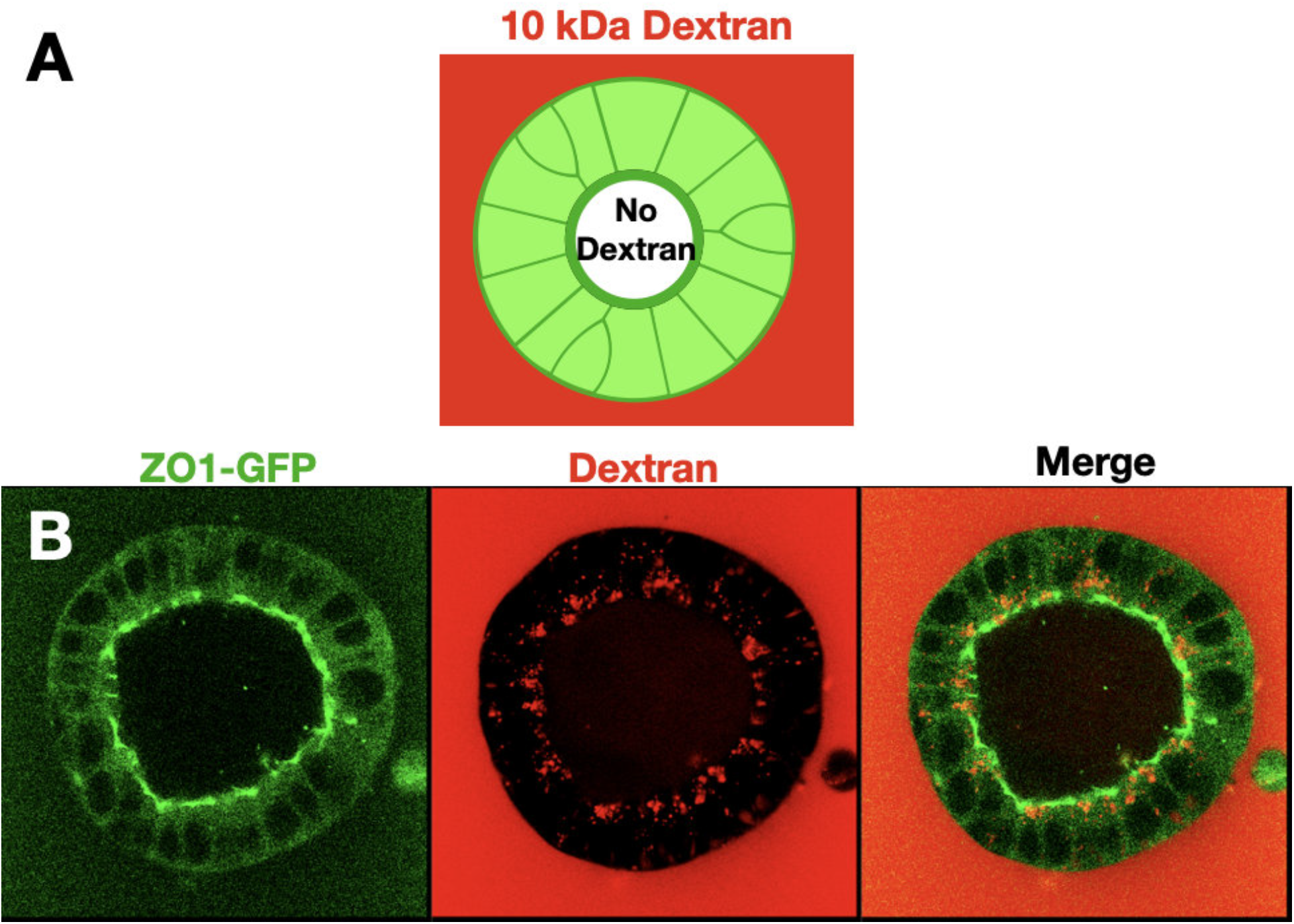
Three-dimensional hPSC form a molecularly isolated niche. (A) Experimental design. hPSC reporter line endogenously expressed tight junction protein ZO1 tagged with GFP (AICS-0023) is used. The 3D culture is exposed to 10kda dextran tagged with Texas Red fluorophore. (B) Horizontal sections show that tight junctions are localized to the inner surface facing the lumen (green). Dextran is visible outside the tissue, but the lumen is devoid of dextran (red). The formation of tight junctions, and the exclusion of Dextran from the inner lumen, suggest that molecules cannot freely diffuse between the media and the inner lumen. We thus conclude that the micropatterned 3D hPSC forms a tightly controlled biochemical niche.

**Fig. S6.**
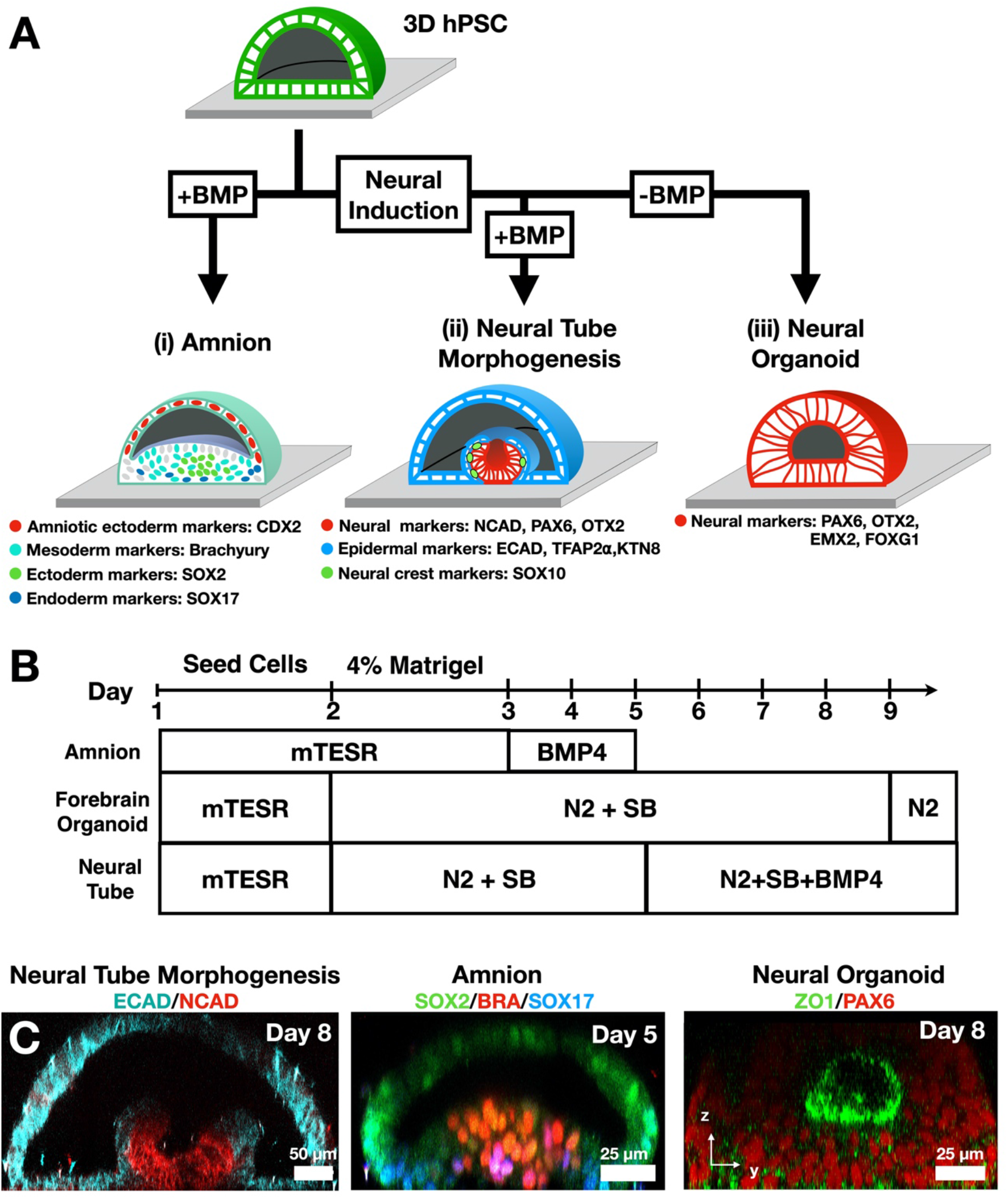
In vitro morphogenesis Road Map. (A) Schematic representation of in vitro morphogenesis in three differentiation protocols. Observed cell markers are indicated, and shown in Figs. S15-16. (i) Exposure to BMP, without neural induction, results in a amnion-like tissue containing cells from three germ layers without formation of a neural fold. (ii) Neural tube morphogenesis is observed when neural induction is followed by exposure to BMP4. (iii) Homogenous expression of forebrain markers is observed under exposure to neural induction without BMP. (B) Experimental timeline for the three protocols. Neural induction media includes N2 supplement and TGFβ inhibitor SB-431542 (SB). (C) Vertical sections of immunostained samples from the three differentiation protocols.

**Fig. S7.**
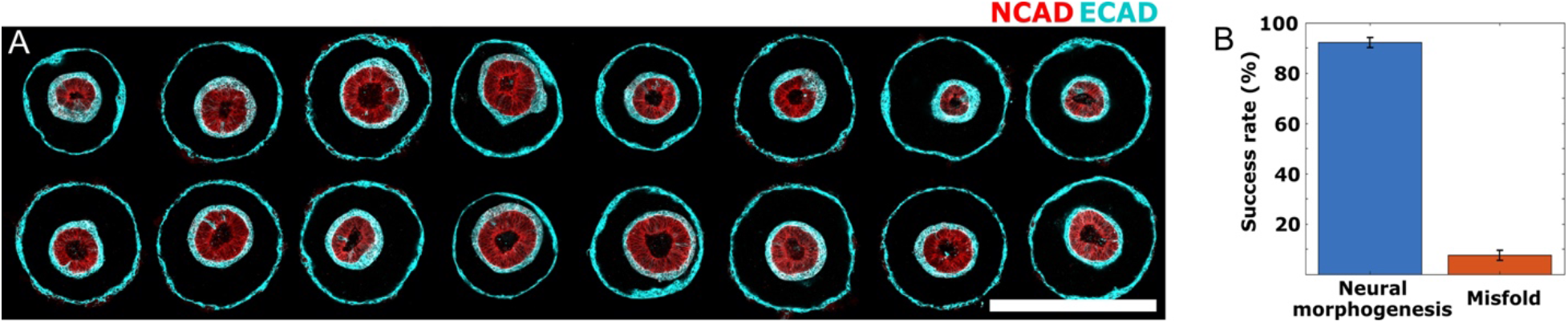
Folding morphogenesis is reproducible. (A) Horizontal sections of 16 circular cultures from a single array exhibit stereotypic fate-patterning and morphology. (B) Bar plot showing success rate of neural tissue folding morphogenesis, in 13 experiments, with total N=100 samples.

**Fig. S8.**
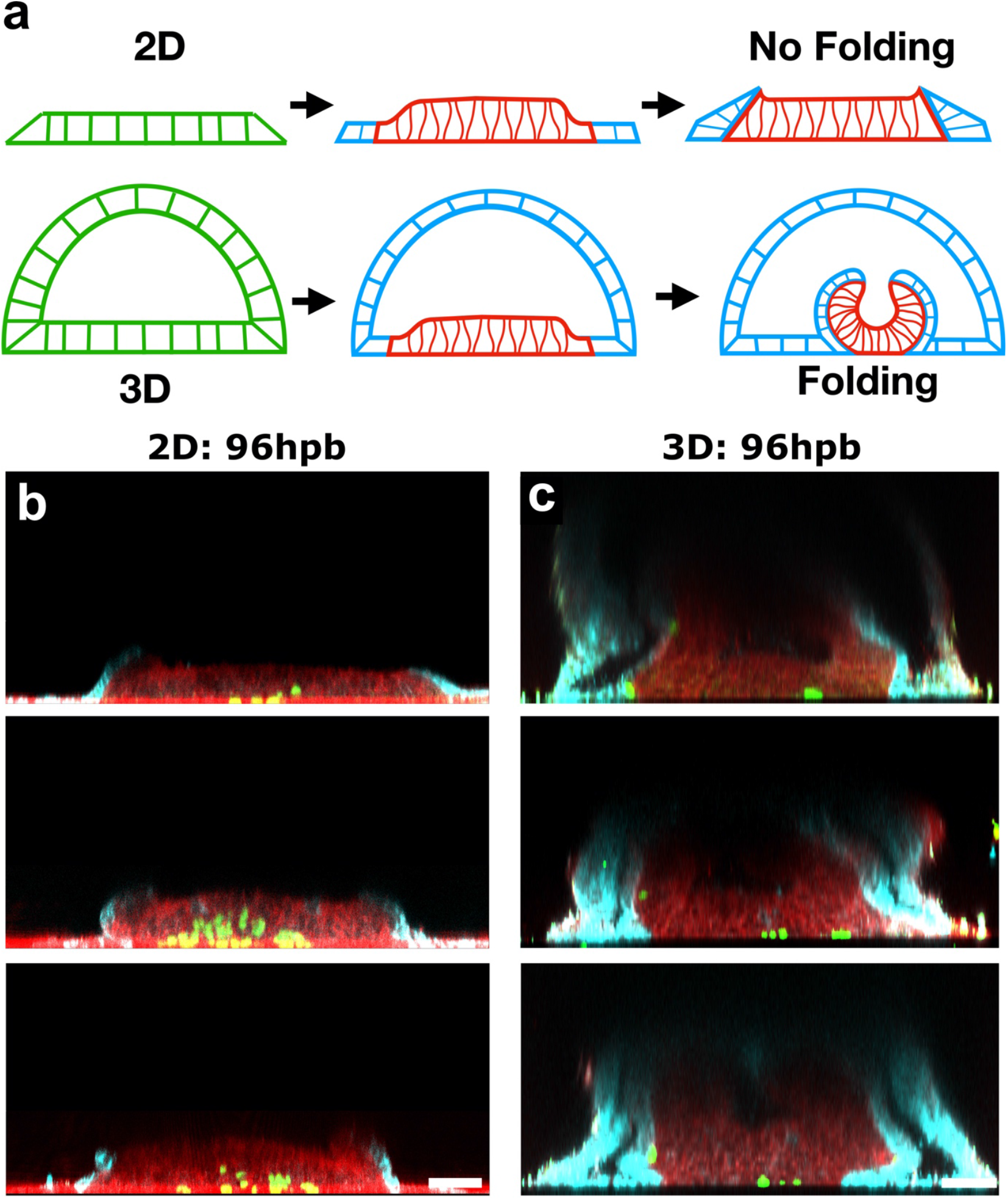
Folding morphogenesis is not observed in 2D micropatterned cultures. (a) Scheme showing experimental design and results. Induction of neural pattern formation in 3D stem cell cultures with a lumen triggers folding morphogenesis. In contrast triggering pattern formation in 2D micropatterned cultures does not trigger folding morphogenesis. (b) Vertical sections of 2D immunostained samples and (b) 3D immunostained samples. NCAD is shown in red, ECAD in cyan, and SOX10 in green. Scale bar is 50μm.

**Fig. S9.**
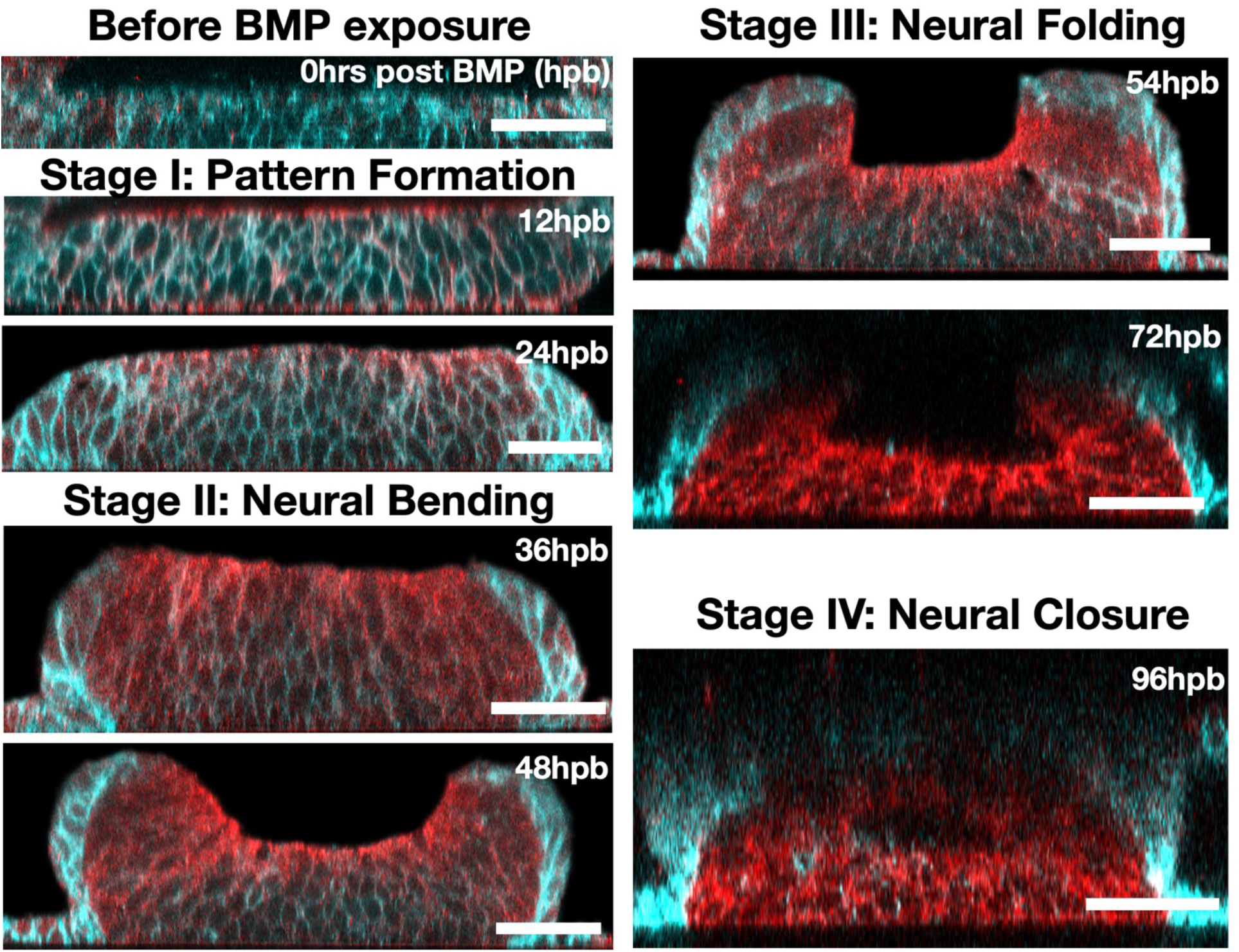
Stages of folding morphogenesis. Vertical sections of immunostained samples during neural fold morphogenesis. ECAD is shown in cyan and NCAD in red. Hours post exposure to BMP (hpb) is indicated. Scale bar is 50μm.

**Fig. S10.**
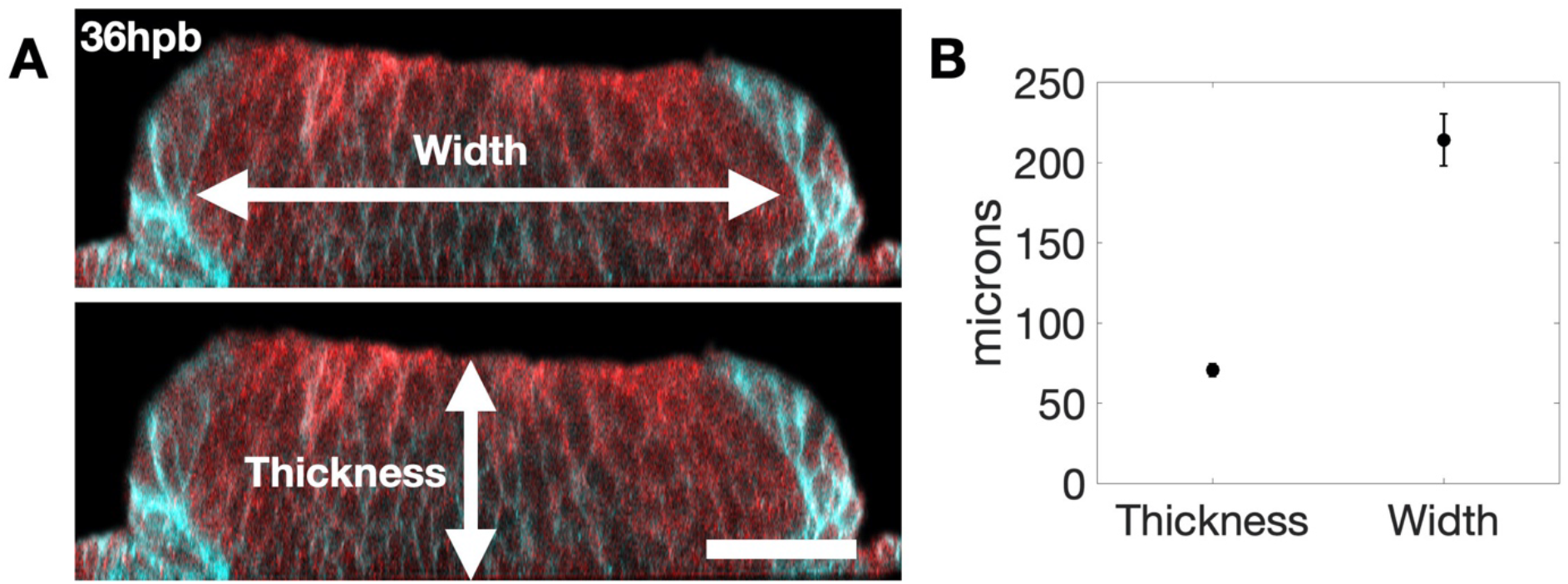
Quantification of neural tissue thickness and width. (A) Vertical section at 36hrs post BMP, and before folding. Scale bar is 50 microns. The neural tissue thickness and width are indicated. (B) Quantification of neural tissue thickness and width averaged over three samples. The average thickness is 70 microns and average width is 215 microns.

**Fig. S11.**
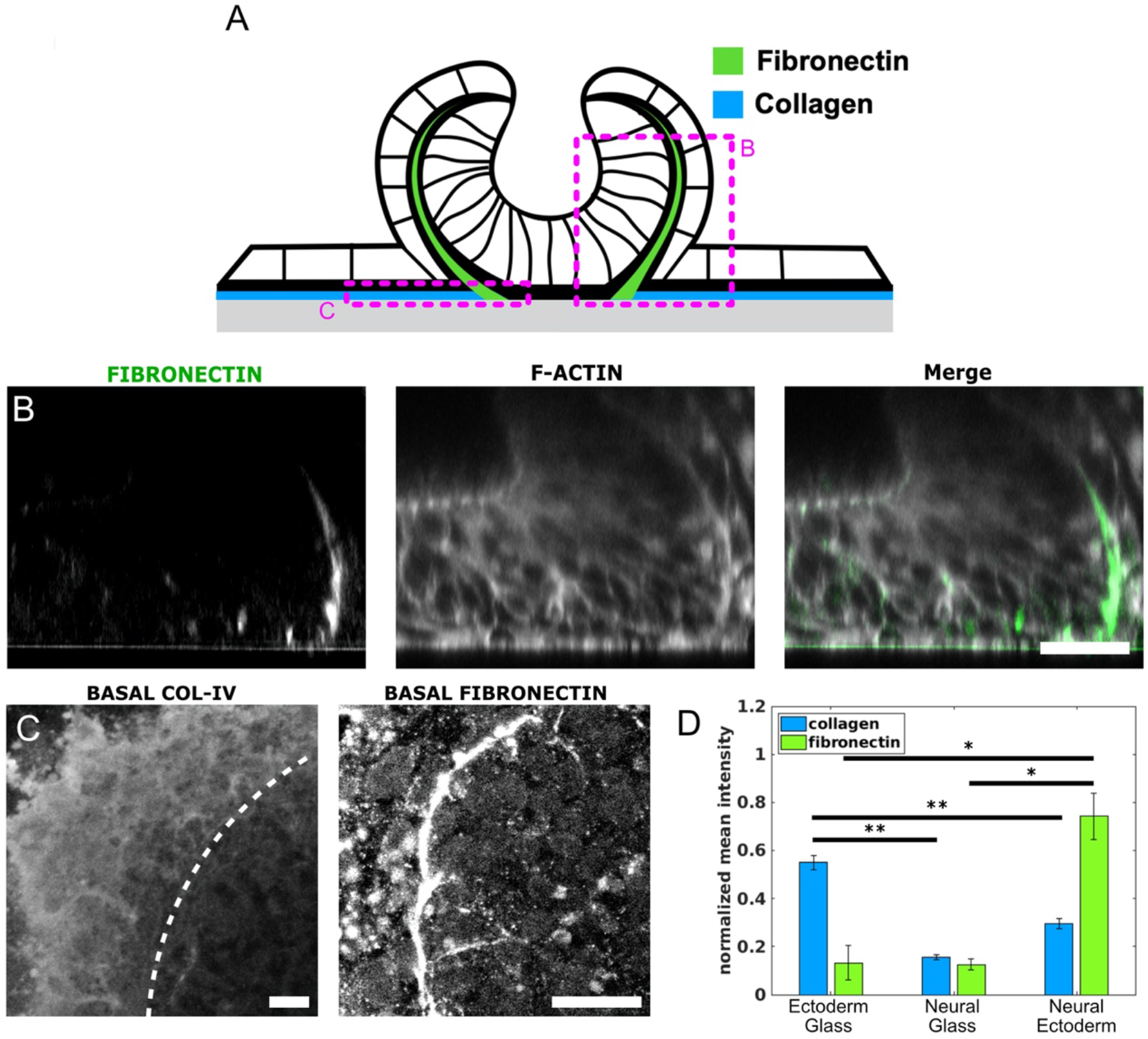
Extracellular composition during neural folding. (A) Scheme showing ECM composition in three interfaces: ectoderm-glass, ectoderm-neural and neural-glass.(B) Vertical sections show fibronectin. (C) Horizontal sections near the glass interface show collagen V and fibronectin.(D) Collagen and fibronectin fluorescence intensity compared at the three interfaces. Scale bars are 25 microns.

**Fig. S12.**
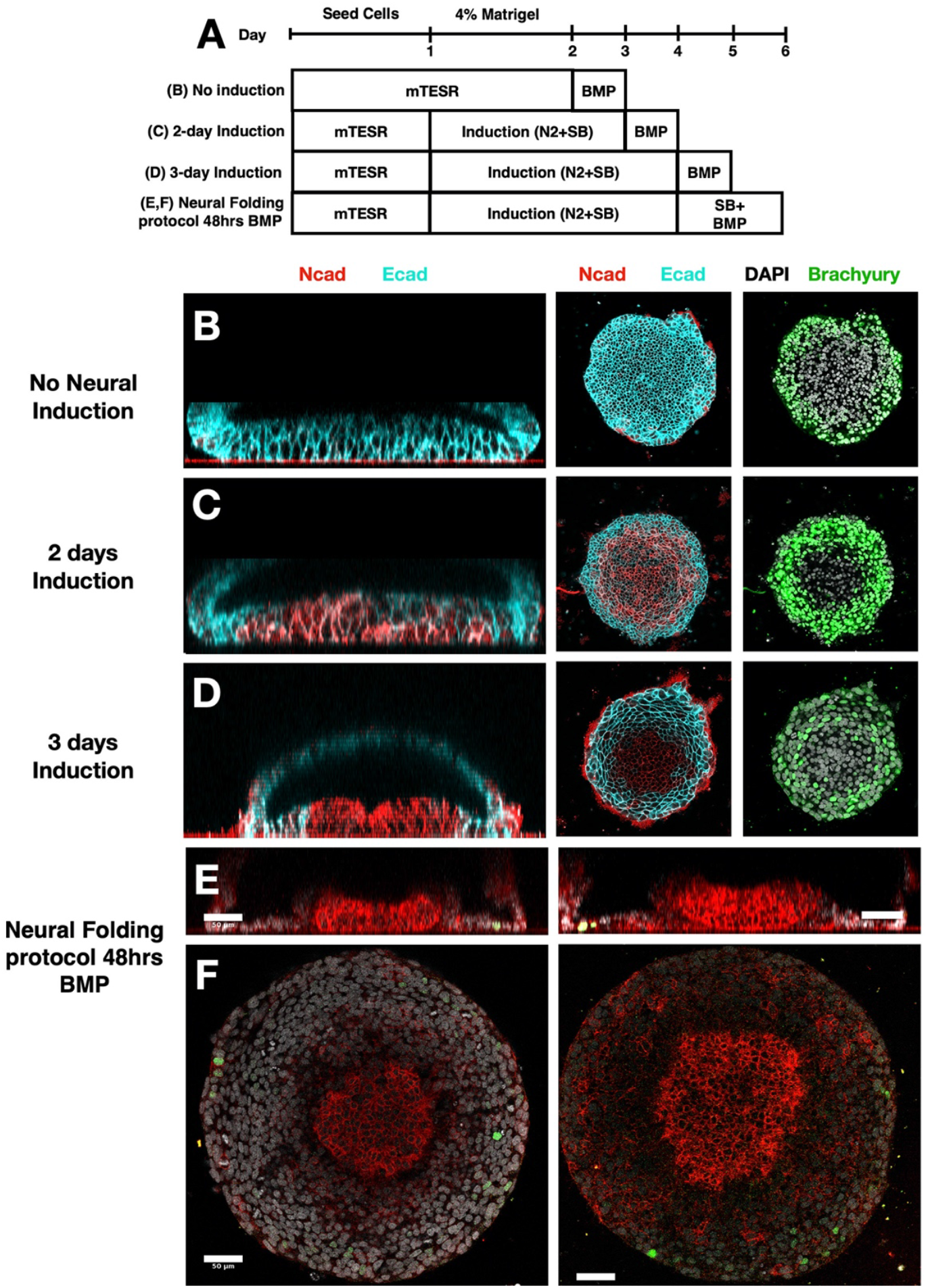
Folding morphogenesis occurs in the absence of mesendoderm tissue. (A) Experimental time line to examine effect of neural induction on cell fate and folding morphogenesis. (B-D) Vertical and horizontal sections showing the neural fates (NCAD) are upregulated with longer neural induction, whereas mesendoderm fates (Brachyury) are downregulated. (E,F) A small number of Brachyury+ cells (<10) is present in the protocol used to for neural tube morphogenesis. Total number of cells in the tissue is ~5000 cells. Scale bar is 50μm.

**Fig. S13.**
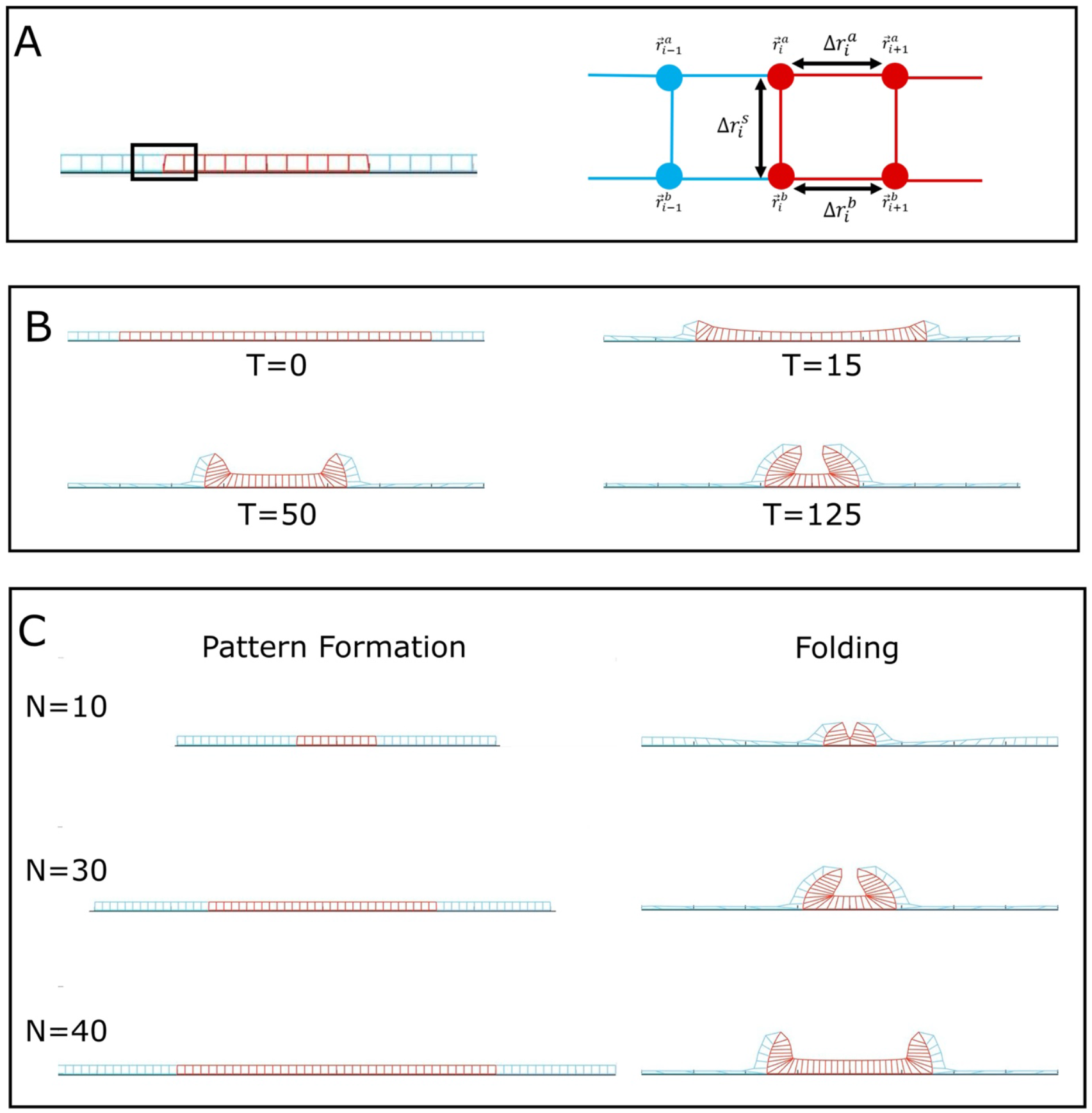
Computational model of neural folding. (A) Zoom-in schematic of vertex model, Δ*r_i_* represents the length of a given springs connecting the vertices. 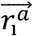 represents the coordinates (x and z) of the given vertex. Here the superscript *a* is for apical points while *b* is for basal points. (B) Time sequence of simulation with 30 neural cells, *m* = 7.5, and μ = 7.5. *m* represents the myosin contractility, while μ is the interface energy per number of cells. Each time unit corresponds to 3000 steps in the simulation. Note that the epidermis stretches and the neural tissue increases its thickness as the simulation progresses, similar to our experiments. (C) The number of neural cells controls the final shape of the folding. Here we show simulations with N=10, 30 and 40 neural cells, while the number of ectoderm cells remains fixed. In the 10 cell configuration we observe the formation of only one central hinge point. In the 30/40 cell configurations two hinge points are formed. The observed tissue shapes match our experimental data.

